# Sounds of silence: electric mobility promises a quieter soundscape for wildlife, but may challenge ultrasonically sensitive species globally

**DOI:** 10.1101/2025.07.10.664227

**Authors:** Baheerathan Murugavel, Manjari Jain, Oliver Lindecke

**Affiliations:** Department of Biological Sciences, Indian Institute of Science Education and Research (IISER) Mohali, Punjab, India; Institute for Biology and Environmental Sciences, Carl-von-Ossietzky Universität Oldenburg; Oldenburg, 26111, Germany

**Keywords:** acoustic orientation, bats, conservation, electric vehicles, e-mobility, green-green dilemma, noise pollution, sensory ecology, ultrasonic noise, urban ecology

## Abstract

Expanding transportation networks generate anthropogenic noise, a stressor for wildlife. The rapid transition to electric mobility, with projections of 40-45 million new electric vehicles (EVs) annually by 2030 (two in five new light-duty vehicles), promises quieter audible environments. However, a critical gap exists in our understanding of the acoustic trade-offs, especially of vehicle-generated ultrasound (frequencies >20 kHz). While ultrasound is known to be biologically potent, used for instance in wildlife repellents, its presence and potential impacts as an emission from traffic on sensitive species, such as bats, remain unaddressed. While the shift to EVs may reduce overall audible noise emissions, offering a green alternative, potential challenges for species with auditory systems tuned to ultrasound, and also actively using it via echolocation, could offset these benefits. This may set up a green-green dilemma, as the objective of decarbonising transport via e-mobility may undermine efforts to restore urban biodiversity. Our systematic review of 49 studies on anthropogenic/vehicular noise impacts on 114 bat species/species groups revealed that road noise, especially when overlapping with bat echolocation and communication frequencies, consistently elicited negative responses in playback experiments, whereas field observations were more variable. A meta-analysis from this dataset showed that the odds of bats being affected significantly increased with both noise intensity and frequency. Building on this evidence, this review explores the implications for a future dominated by EVs. We highlight the urgent need for further research to define acoustic thresholds and develop conservation strategies for wildlife in rapidly urbanising and electrifying landscapes.

## Introduction

Human-induced anthropogenic stressors have fundamentally modified natural ecosystems leading to the dawn of the Anthropocene, a new geological epoch (Monastersky 2015). In this rapidly changing environment, wildlife is increasingly exposed to novel conditions, such as increased noise and light levels at unprecedented spatiotemporal scales (Sanders et al. 2021). A marked consequence of urban expansion is the growth of transportation networks, leading to a rise in land and air traffic (Coffin 2007). These developments modify acoustic environments, often elevating noise levels in previously quiet areas. Such noise pollution may alter movement behaviour and energy expenditure (Oliveira et al. 2021; Byrne et al. 2022), affect metabolism and food intake (Song et al. 2020), as well as mask or distract from critical biological signals necessary for communication (Leonard and Horn 2012; Stelling-Kończak et al. 2016; Legett et al. 2020; Vieira et al. 2021; Ye et al. 2025), foraging (Siemers and Schaub 2011; Mason et al. 2016; Blair et al. 2016; Tennessen et al. 2024) and reproduction (Schroeder et al. 2012; Senzaki et al. 2018), thereby undermining population viability and altering community dynamics (Bunkley et al. 2017; Iglesias-Merchan et al. 2018; Senzaki et al. 2020, 2023), ultimately posing significant threats to biodiversity. However, such changes typically occur at rates that exceed the life spans of long-lived organisms, potentially disrupting community structures as the most adaptable organisms outcompete less flexible ones in rapidly changing environments (Duquette et al. 2021).

Anthropogenic changes do not universally harm wildlife; some species may adapt or even thrive under novel conditions. For instance, several species of birds and bats are called ‘urban exploiters’ (Kark et al. 2007; Møller and Díaz 2018). Urban wildlife cope with altered soundscapes by either reducing vocalisations – a phenomenon called the *silentium* effect (Sun and Narins 2005; Pearson and Clarke 2019) – or by increasing the call amplitude, the Lombard effect (Brumm and Zollinger 2011; Hotchkin and Parks 2013). Cities also provide vacant niches that adaptable species can exploit (Evans et al. 2009), thereby increasing urban biodiversity, a process referred to as “biological urbanisation” (Moller 2014). The colonisation may be driven by *push* factors such as habitat loss in surrounding landscapes, i.e. natural habitats, or by *pull* factors such as abundant forage from exotic plants and reduced competition, as well as predation pressure (Moll et al. 2018; Eötvös et al. 2018; Lill 2020).

Yet animals’ freedom to move remains constrained by transportation infrastructure, not only through physical impediments but, crucially, also through sensory disruption (Coffin 2007; Bednarz 2021). In bats, for example, artificial illumination has been shown to alter movement behaviour and route choice (Voigt et al. 2018; Barré et al. 2020). By analogy, the broadband noise that accompanies road networks can function as an acoustic barrier to bats, in rural (Schaub et al. 2008; Berthinussen and Altringham 2012; Bhardwaj et al. 2021; Ye et al. 2025), as well as urban areas (Lehrer et al. 2021). Nonetheless, many cities harbor a surprisingly rich bat diversity, implying rapid behavioural plasticity under chronically elevated noise regimes (Voigt and Kingston 2016).

Roughly 90% of extant bat species orient and forage with an echolocation biosonar, a highly specialised sensory modality involving the emission of ultrasonic pulses and subsequent interpretation of returning echoes to create detailed acoustic images of their surroundings (Altringham et al. 1996). When anthropogenic noise stays within the dynamic and spectral limits of a bat’s auditory filters, individuals can, in principle, compensate by shifting call frequency, timing or amplitude, allowing them to persist in noisy environments. This response would increase the adaptability of bats, resulting in urban-adaptive species (Luniak 2004). However, noise that exceeds those limits of sensory processing imposes energetic and informational costs that drive local exclusion and ultimately erode community diversity in such environments. Ultrasound noise specifically has been negatively affecting a wide range of taxa and is often used as a deterrent for rodents, dogs, deer and bats (Blackshaw et al. 1990; Arnett et al. 2013; Aflitto and Hofstetter 2014; Laguna et al. 2022)

What we still do not know is where the boundary lies between tolerable and exclusionary noise for most species, or whether the soundscapes of future transport networks will cross that boundary. Despite a growing literature on current (conventional) traffic noise effects on animals, systematic investigations that map noise spectra of next-generation vehicles in urban landscapes onto species-specific hearing and processing limits are virtually absent, leaving planners without the evidence needed for wildlife-friendly road design and vehicle engineering. However, global sales of new electric vehicles (EVs) are on a steep upward trajectory, with multiple leading energy and market analysis groups projecting a dramatic increase by 2030 (I.E.A. 2024). Forecasts indicate that EVs could constitute anywhere from nearly one-third to over two-thirds of all new passenger cars sold worldwide by the end of the decade, translating to tens of millions of units annually (Maffei and Masullo 2014; Sen and Miller 2023). Notably, hybrid electric vehicles (HEV) that combine a conventional internal combustion engine and electric engines for propulsion add to potential ultrasound emissions.

The unprecedented global shift toward electric mobility offers an unexpected conservation lever. Battery-electric drivetrains eliminate the combustion-engine roar that dominates the 0.1 to 10 kHz band (Maffei and Masullo 2014; Praticò et al. 2020), thereby lowering sound-pressure levels most salient to humans and many terrestrial vertebrates. Pilot monitoring programmes already report reductions of 15 to 20 dB at idle relative to comparable internal-combustion vehicles (ICVs), with similar gains during low-speed deceleration (Sandberg et al. 2010; Lan et al. 2023). By diminishing this sonic masking layer, e-mobility could create *de facto* acoustic refugia along existing road networks and facilitate recolonisation by acoustically sensitive taxa (Yosef et al. 2021). From existing knowledge on how present traffic noise, especially on roadways, impacts various species, it is high time to look into the future to understand and predict how acoustically dependent wildlife will respond if the existing road networks emit less deterring noise.

### A green-green dilemma invoked by unregulated ultrasound?

Paradoxically, the same technology that quiets the (human) audible spectrum may introduce a new pollutant in the ultrasonic domain. While EVs and hybrid-electric vehicles (HEVs) reduce many types of audible noise (particularly engine-related), the possibility of increased ultrasonic emissions generated from several EV components, such as high-frequency inverters and power electronics, is not negligible (Staneva et al. 2020; Fig.1). Concerns have been raised in the engineering research community about significant gaps still existing in environmental assessments of EVs, including their emission spectra (Hawkins et al. 2012) but these do not echo much in the wildlife research community. However, converting DC current from the battery to AC for the motor often involves electronic switching at high frequencies, some of which can extend into the ultrasonic range. Furthermore, auxiliary systems, i.e. components such as DC-DC converters, battery management systems, magnetostrictive motor laminations, and certain sensors, can emit ultrasonic vibrations. Although the intensity and ecological relevance of these emissions have not been thoroughly quantified, first reports suggest the presence of measurable ultrasound emissions. Above all, high-speed cooling fans and field measurements at urban charging hubs document supraharmonic noise spanning 2 to 150 kHz, with prominent peaks at 22 kHz and 40 kHz (Schöttke et al. 2014; González-Ramos et al. 2024; Clar-Garcia et al. 2025), i.e., frequencies overlapping the echolocation and social-call spectra of many species of bat (Fenton and Bell 1981) as well as the hearing ranges of diverse insects (Nakano et al. 2015; Geipel et al. 2021; Barber et al. 2022) and mammals, e.g., rodents and carnivores, including domestic dogs (Heffner and Heffner 1985, 2007; Powell and Zielinski 1989; Blackshaw et al. 1990; Ramsier et al. 2012).

**Fig. 1.**
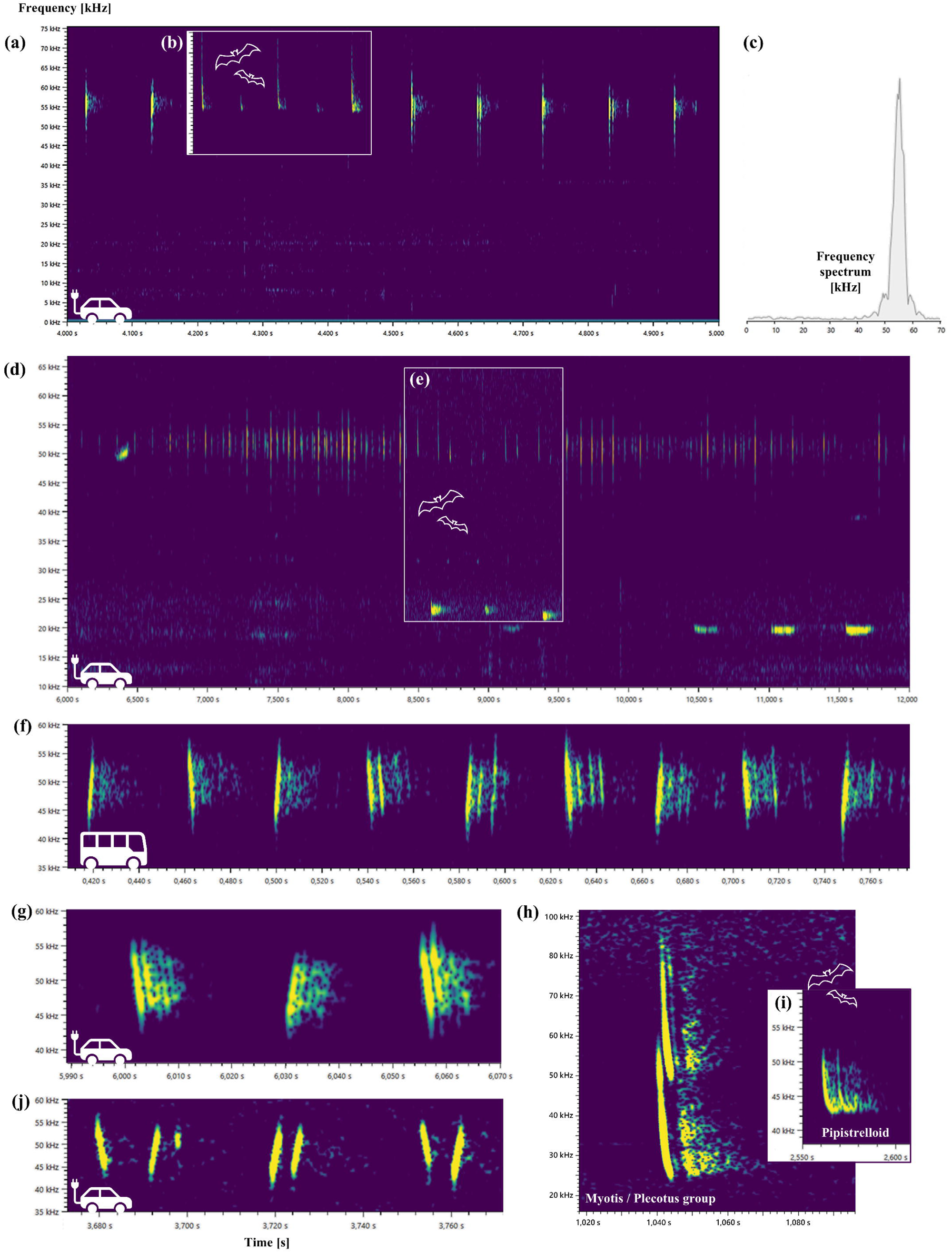
Spectrograms of electric vehicles randomly sampled in city traffic match bats’ dominant frequencies and pulse spacing, highlighting substantial masking potential. (a) A one-second snippet of a car (Tesla Model X) passing by in a 30-km/h zone, incl. an inlet showing a bat calling sequence. The car’s acoustic noise is biased toward the ultrasonic frequency range (c), inaudible to humans, and shares characteristics with bat calls (b). The 40 to 60 kHz spectrum is typical for e-vehicles (a; d; f, public e-bus; g, Škoda Elroq; j, BYD Atto 3). Recordings of a VW id.3 in a 50-km/h zone show additional emissions at 20 kHz (d), which are typical for echolocation calls of open-space foraging bats. The inlet (e) shows calls of two species of bats. Examples of bat groups (h, i) that are commonly misidentified in automatic analyses and on audio/visual-based bat detectors.

The current policy and regulatory frameworks focusing on vehicle acoustic emissions are largely centred around audible noise. With EVs, also mandatory pedestrian alert sounds, so-called ‘acoustic vehicle alerting systems’ (often artificial and tonal) are imposed at low speeds to ensure safety (Hastings et al. 2011; In this work, we are not further addressing these alert systems) . However, these regulations rarely address ultrasonic frequencies. As EV adoption increases, it is crucial to investigate whether their ultrasonic emissions have any significant ecological impacts, especially on wildlife species sensitive to high-frequency sound.

This phenomenon sets up a classic green-green dilemma: one environmental objective, namely decarbonising transport through widespread EV adoption, may undermine another, namely the restoration of biodiversity in anthropogenic landscapes. A conflict between two seemingly positive environmental goals in the context of renewable energy and biodiversity conservation emerges, dividing involved parties and provoking emotional responses in involved groups that each work towards a sustainability goal (Voigt et al. 2019). However, a growing recognition of “ultrasound noise pollution” in cities has prompted the first calls for regulatory thresholds analogous to those that govern audible noise (Grimshaw-Aagaard and Bemman 2024).

Mitigating this emerging conflict will require (i) systematic mapping of acoustic sensitivity of wildlife with a focus on EV-related ultrasonic emissions potentially emerging in traffic scenarios (ii) incorporation of ultrasonic bands into vehicle and infrastructure noise standards, and (iii) engineering solutions, such as acoustic damping or shifting switching frequencies above 150 kHz, to push tonal artefacts beyond the biologically relevant range, i.e. animal hearing and echolocations thresholds. Until such measures are implemented, the putative acoustic benefits of e-mobility remain contingent on a technology that may, inadvertently, export noise into a spectral window critical for non-human species.

Here, using bats as a model, we assess the sensitivity of a globally occurring animal group to current traffic presence, i.e., its noise. Through a systematic analysis of studies, we aim to identify commonalities, the acoustic dimensions that appear problematic, and whether anthropogenic ultrasound emissions have been adequately addressed to date. Specifically, we plan to first quantify the number of bat species affected by different types of anthropogenic/vehicular noise globally. Next, we aim to assess how bat species are likely to be influenced by the acoustic characteristics of noise, mainly the intensity and frequency. We additionally compare and discuss the implications of the findings in the context of future soundscapes and noise expected to be generated by EVs in comparison with typical bat call emissions.

## Methods

### Literature review and database search

We followed the Preferred Reporting Items for Systematic reviews and Meta-Analyses (PRISMA) guidelines for ecology and evolution while reviewing and refining the literature for this study (Moher et al. 2009; O’Dea et al. 2021). Terms “bats”, “traffic” and “noise” were used to search literature in the Web of Science database. Searches were first done only on the Web of Science Core Collection. Next, searches were conducted on all databases including the Web of Science Core Collection, Grants Index, KCI-Korean Journal Database, Preprint Citation Index, ProQuest™ Dissertations & Theses Citation Index, and SciELO Citation Index Databases. These databases are also sensitive to Boolean operators so searches were conducted for the “AND” operator to see how the results differ. Search results were exported in RIS and BibTex format. All searches were done on the 1^st^ of April 2025, which before screening yielded studies from the year 1956 – 2025.

### Deduplication and primary screening

Removal of repeated results from different search queries (deduplication) is another key step that needs to be described well in a reproducibility context. EndNote online reference management tool with the ‘remove duplicates’ option was used to eliminate duplicates from all the Web of Science results, followed by manual deduplication of entries using Zotero offline reference management tool. The deduplicated entries were manually screened for relevance to our questions based on titles. Irrelevant entries such as brown adipose tissues, blunt abdominal trauma (abbreviated to BATs), traffic and Artificial Intelligence literature, etc were eliminated. At this stage, the relevant studies covered the years 1997 - 2025

### Abstract screening and final dataset

Since the effect of anthropogenic noise on bats is the primary objective of this study, the abstracts and full texts of the 146 studies were screened, and 49 relevant ones were found to have investigated the effects of anthropogenic noise on bats. Reviews, some basic sensory ecology studies that had used broadband filtered white noise, reports of road kill, etc. that did not investigate effects of anthropogenic noise on bats were removed from the final dataset (Supplementary Fig. 1).

### Information collected from studies

From the shortlisted studies, firstly, the list of species studied was extracted. Next, whether anthropogenic noise (hereafter noise) was reported or not was noted as a binary outcome. For studies with noise information available, details on the species studied, location (country level), reported sound intensity, frequency of noise, type of noise, study design, and effect of noise/traffic were extracted at a species level. Families of each species reported were included for further comparisons. This was obtained directly from the studies, if reported or collected from the Encyclopedia of Life traitbank (http://eol.org). Noise type was categorised into four major classes, namely Road, Rail, Air, and others (Supplementary Table 1). Some studies had recorded vehicular noise from roads and used it as playbacks in either lab or wild experiments. Such cases where road noise was used but not directly from the roads were classified as Road* (Supplementary Table 1).

### Noise information and effects on bats

Sound intensities of anthropogenic/traffic noise that were reported/tested on bats were extracted from the studies and were classified into 11 categories of 0 to >100 decibels (Supplementary Table 2). Frequency information, whenever reported, was extracted and categorised similarly to intensity. Frequency levels ranged from 0-100 kHz, and they were categorised into 10 categories (Supplementary Table 2). However, not all studies that report the noise frequencies use that in their analysis, as they either choose a frequency that does not overlap with bat calling frequencies (Domer et al. 2021) or mention that they exclude frequency from analysis (Cory-Toussaint 2022).

From the results of the studies with and without noise, based on the description of the studies, effects were first scored as positive, negative, no effect, neutral effect, or not applicable (refer to Supplementary Table 2 for details). However, for further analysis, only studies with noise information available were used. Here, effects were classified as a binary outcome (affected or not affected). Studies that had reported that noise treatments/conditions had ‘no effect’ on a species were scored as ‘not affected’, and all other effects as ‘affected’ by noise.

### Statistical analysis

All analysis were done in R (R Core Team 2021). All the species that were exposed to anthropogenic noise were selected, and the effects of sound intensity and frequency were compared. With a binary response variable of noise effect (affected = 1 and not affected = 0), we fitted two Generalised Linear Models (glms) with a binomial error distribution and a logit link function to test whether the effect of noise increased with an increase in intensity (model1) and frequency (model2). We used the previously assigned categories for both intensity and frequency (n=11 with 10 dB and n=10 with 10 kHz bins, respectively). Models were run using the ‘glm’ function in the *stats* package, and data handling was done using the *dplyr* package.

## Results

### Literature trends

A total of 4951 entries were obtained from the database searches. The deduplication step resulted in a dataset of 1362 entries, which, on further manual screening finally provided 49 studies that had studied the effects of either anthropogenic noise or traffic conditions or both on bats (Supplementary Fig. 1). Globally, this was studied in 21 countries, of which 16 were High-Income countries according to the World Bank classification (https://datahelpdesk.worldbank.org/; Fig. 2a). With a few exceptions (Geipel et al. 2019; Domer et al. 2021; Cory-Toussaint and Taylor 2022), no studies have been conducted in Central America, most of Africa, Western, Central, Southern and Southeastern Asia on this topic yet.

**Fig. 2.**
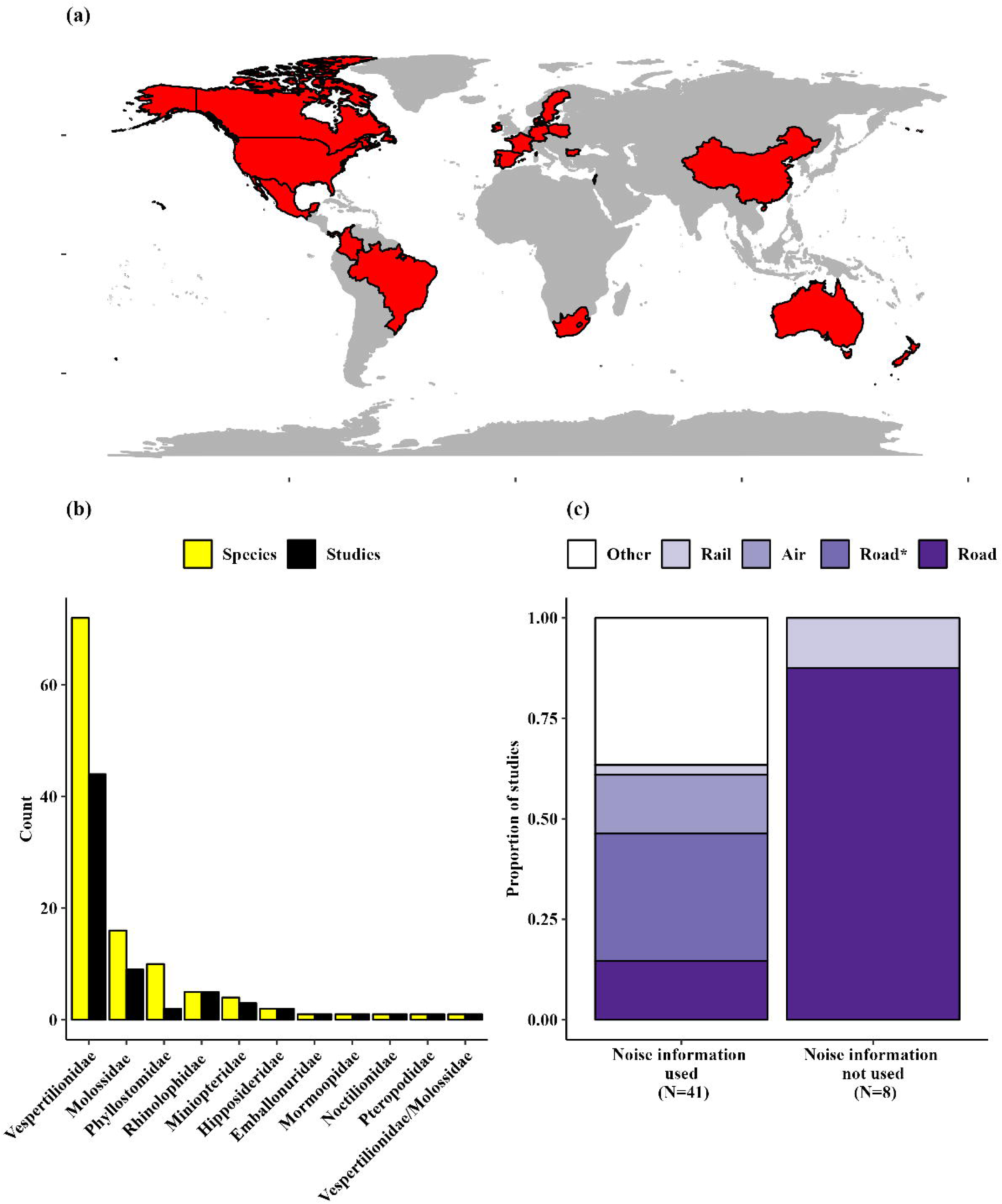
Summary of literature review on effects of anthropogenic noise and traffic on bats with (a) countries studied in red (b) number of studies and species reported and (c) types of noise used, where N denotes the number of studies.

A total of 114 species or species groups were studied that comprised 10 bat families. Family Vespertilionidae was the most studied (N=72 species), followed by Mollossidae (N=16 species), and families Emballonuridae, Mormoopidae, Noctilionidae, and Pteropodidae were the least studied (N=1 species each) and no studies have been conducted on species from 11 bat families, some of them often occupying anthropogenic landscapes of the world (Fig. 2b). Out of the noise types, road noise was most explored (N=26 studies), followed by other noises (N=15 studies), including environmental soundscapes, motorcycle rallies, music festivals, gas or compressor stations and acoustic deterrents. Rail noise was the least studied, followed by air noises (Fig. 2c; details in Supplementary Table 1). It should be noted that not all studies conducted with road noise exposed bats to direct road traffic. Some used recorded traffic noise and played it back for experiments conducted in controlled lab conditions or the wild (N= 13 studies; Supplementary Table 1).

### Effect of noise

Amongst the 114 studied species, 101 were exposed to anthropogenic/vehicular noise out of which an effect was reported in 46 species whereas 78 species were not affected by the noise treatments/conditions and contexts studied (Fig. 3a). Intraspecific differences in effects were found and was due to differences in noise type, study locations and experimental durations. The same species showing an effect in contexts such as road noise, proximity to roads, playback noise etc were not affected in other contexts such as air traffic, road types, and compressor stations. In some cases, species showed an effect in some locations but not in others (refer to Supplementary Table 2 for details). While the majority of the species were negatively affected, there were four cases where noise treatment had an effect but could not be classified as negative (Fig. 3a). All species that were subjected to road noise in playback either in the lab or in the field experiments showed were affected by noise treatments, but studies that evaluated the effects directly around roads showed an effect in only 32% of the cases (Fig. 3b). However, all the species exposed to rail noise were negatively affected, and air noise affected bats in 84% of the cases. Noises from other sources such as compressor stations, music festivals, acoustic deterrents and motorcycle rallies had an impact in 30% of the cases (Fig. 3b)

a. *Intensity* Anthropogenic noise spanned across intensities ranging from 10 to >100 dB (SPL) in the studies. One study using drones reported an exposure of 0.5 dB noise relative to their noise floor used and was excluded from this analysis. All categories above 20 dB affected the bats (Supplementary Fig. 2a), but depending on the contexts and conditions, some species were not affected by the intensity levels exposed (even above 20 dB). The logistic regression (model 1) predicted a significant positive relationship between sound intensity and the likelihood of species getting affected (β = 0.241 ± 0.055 SE, *z* = 4.40, *p* < 0.001; Supplementary Fig. 2b). In other words, species exposed to higher sound intensities were significantly more likely to be affected. For every 10 dB increase in noise levels, the odds of species getting affected increased by 27.3%. Overall, 46 species were affected by either one or more sound intensity categories, with *Lasionycteris noctivagans*, a species of the western Palaearctic region, being the most affected species that showed an effect from 20 to 100 dB (SPL) (Supplementary Fig. 3a)
b. *Frequency* Noise frequency information was available for 74 species overall and ranged from 0 to 100 kHz (Supplementary Table 2). Forty (54%) of these species were affected by the noise treatments (Fig. 4a). All studies that had used frequencies above 30 kHz showed an effect on the bats and seven species/species groups were affected by all the 10 frequency categories (Supplementary Fig. 3b). Some studies had reported frequencies but had not used them in their analysis, i.e, the observed effect of noise might not be directly attributed to an effect of frequency used. However, we included such studies for our analysis. Logistic regression (model 2) also predicted a significant positive relationship between noise frequency and the likelihood of species getting affected (β = 0.84, SE = 0.17, z = 4.94, p < 0.001). Every 10 kHz increase in noise frequency increased the odds of species getting affected by 132 % (Fig. 4b).

**Fig. 3.**
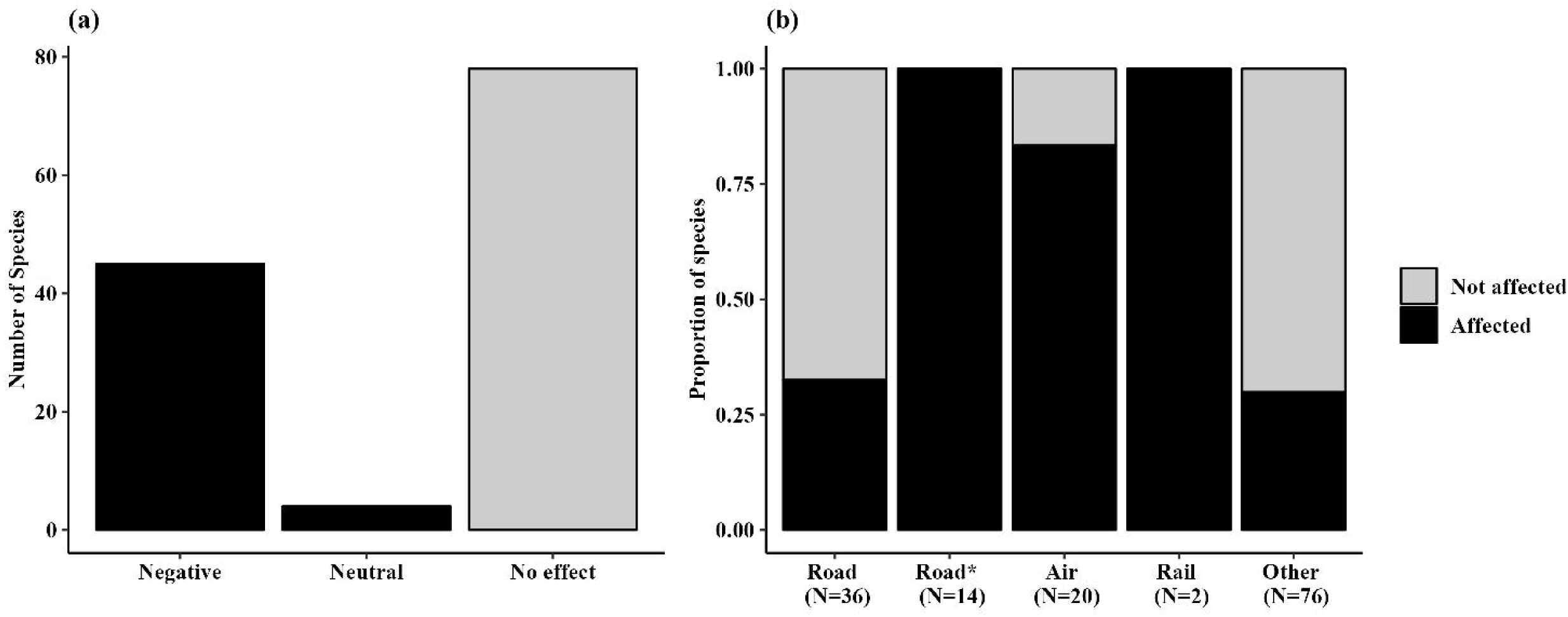
Effect of anthropogenic/vehicular noise on bat species. (a) Species counts relative to the type of effects reported in the studies that exposed bats to noise. (b) Proportion of species affected by different types of noise exposures [* indicates species that were exposed to road noise indirectly via playback experiments either in the wild or under lab conditions, and N denotes the number of species exposed]

**Fig. 4.**
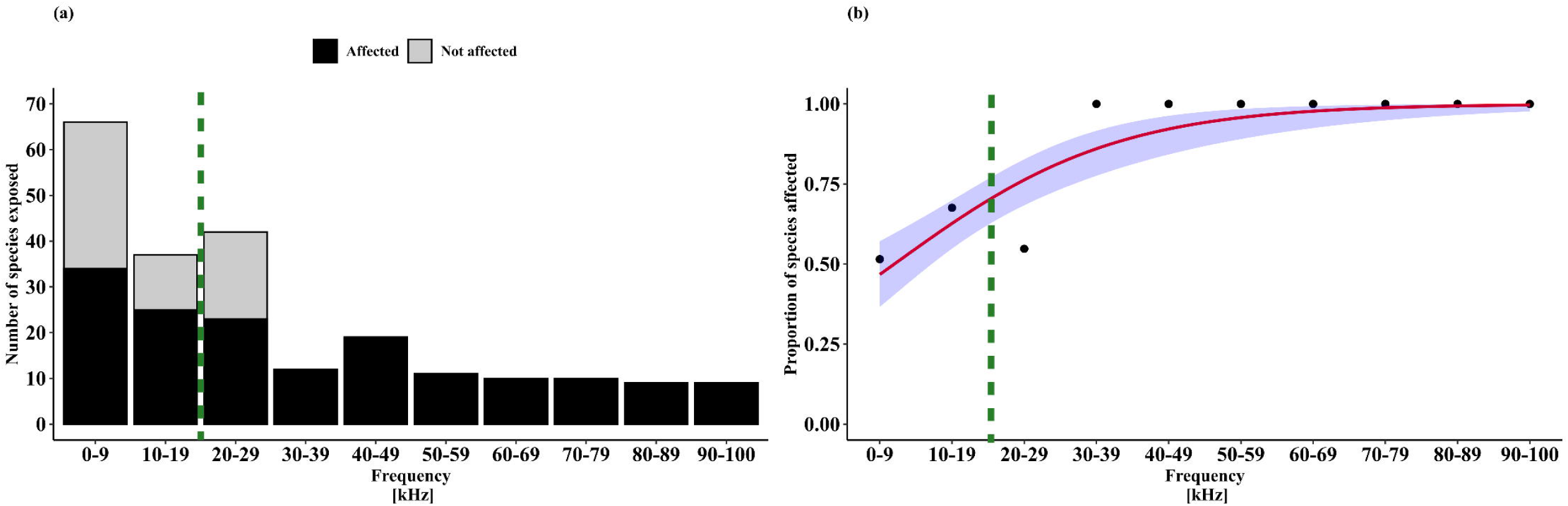
(a) Number and (b) proportion of species/species groups exposed and affected by noise frequency categories. The predicted probabilities from the linear regression model are plotted as red lines with a 95% confidence (blue) ribbons. Green dashed lines denote the ultrasound border, and a clear effect with 100% probability is seen in all species/species groups exposed to frequencies >30 kHz, suggesting a potential impact of ultrasound noise in bats.

## Discussion

Our systematic review aimed to establish the acoustic impact of traffic noise on bats and identify thresholds by synthesising current knowledge on both audible and ultrasonic pollution, to support an evaluation of the wildlife-friendliness of the global transition to electric mobility. It delivers three answers to knowledge gaps that we articulated *a priori*: (i) where the transition from ‘tolerable’ to ‘exclusionary’ noise for bats begins, (ii) which acoustic dimensions are most likely to push species beyond that boundary, and (iii) how the forthcoming dominance of electric mobility could shift these thresholds within urban soundscapes. We confirm that both noise intensity and frequency are significant predictors of negative impacts. Crucially, our meta-analysis shows that the probability of a detectable effect rises steeply once sound-pressure levels exceed ∼20 dB SPL (Supplementary Fig. 2) and/or spectral energy overlaps bat call space above ∼30 kHz (Fig. 4). These numerical thresholds begin to delineate the critical “tolerable–exclusionary” boundary for bats.

The congruence of our global synthesis with diverse case studies, which have linked anthropogenic noise to impacts on mating behaviour, foraging efficiency, metabolism, and gene expression (eg. Jiang et al. 2019; Song et al. 2020; Ye et al. 2025), support the inference that acoustic masking is a principal mechanism driving these widespread effects in bats. However, due to the limitations of available studies in aspects of species diversity, behavioural context (e.g. foraging, mating, migration) and regional foci, we cannot yet make species-specific recommendations but we may expect some species could be more vulnerable than others in future soundscapes. For example, traffic noise can potentially mask the ability of bats to detect prey, leading to a direct consequence on their ecology. The prey detection ability of the greater mouse-eared bats, *Myotis myotis,* is affected by background traffic noise under laboratory conditions (Schaub et al. 2008), showing a direct negative impact of noise on this species. It is crucial to consider the ecological impacts of such factors in urban planning, particularly in the context of urban rewilding, an approach aimed at boosting biodiversity in human-dominated areas by improving habitat quality for wildlife (Finnerty et al. 2025). The success of rewilding will depend not only on restoring physical habitats but also on ensuring that acoustic environments support the sensory ecology of target species, rather than undermining it by inadvertently creating ecological traps (Hale and Swearer 2016; Goławski et al. 2025).

Future soundscapes enriched with ultrasound (Fletcher et al. 2018; van Wieringen and Glorieux 2018) therefore need to be accounted for, especially when these are fairly predictable, as in the case of EV emissions, where measurements and modelling are possible anytime based on the present-day EV fleet. Types of noise studied in bats to date highlight impacts across behavioural and physiological scales: Allen et al. (2010) found that bats roosting in urban areas exposed to anthropogenic stressors had larger birth sizes and higher growth rates compared to cave-roosting bats. Chronic exposure to anthropogenic noise has been linked to reduced food intake and altered gene expression, specifically in genes associated with metabolism and immune functions (Song et al. 2020). Behavioural avoidance of noisy areas is reported in many species (Schaub et al. 2008; Borkin et al. 2019; Ellerbrok et al. 2022) incurring direct foraging bats. Overall, these studies demonstrate that heightened noise can affect bat health and behaviour with potential implications for disease dynamics in human-dominated environments.

Exemplary field recordings of EVs and HEVs demonstrate a tangible 4–10 dB reduction in the audible <10 kHz band (Maffei and Masullo 2014; Staneva et al. 2020; Praticò et al. 2020) but tonal peaks beyond 20 kHz are neglected, probably because these are irrelevant where human noise guidelines need to be met (Leighton 2018). Thus, while electrification may indeed create acoustic refugia for taxa limited by low-frequency engine roar (Yosef et al. 2021), it risks crossing the ultrasound threshold we identified for species whose echolocation or communication energy lies above 30 kHz. This confirms the central hypothesis of our study: the solution to one stressor (decarbonising transport) introduces another, a classic green-green dilemma. However, this dilemma is particularly acute because of a historical blind spot our review uncovered: the profound impact of high-frequency noise has been largely overlooked in past traffic studies that focused on bats. This knowledge gap is compounded by significant geographical and taxonomic biases in the literature (Fig. 2a, b). With research heavily concentrated in the Global North and on a single bat family (Vespertilionidae), addressing this emerging conflict between e-mobility and ultrasound-sensitive wildlife therefore, requires a clear path forward, beginning with methodology. We call for a standardised, full-spectrum (0-100 kHz), distance-normalised reporting protocol to overcome the inconsistencies that have hampered past research. Beyond methods, mitigating this dilemma will necessitate (i) systematic mapping of acoustic sensitivity across diverse wildlife, focusing on EV-related emissions; (ii) incorporation of ultrasonic bands into vehicle and infrastructure noise standards; and (iii) engineering solutions to push tonal artefacts beyond the biologically relevant range. The ∼20 dB and ∼30 kHz thresholds derived here provide the first evidence-based, quantitative targets for such engineering and policy efforts.

However, we found a discrepancy between controlled playback experiments (which consistently show effects) and direct field observations (which are more variable). Acoustic parameters such as intensity and frequency appear to generally negatively affect bats when heightened, but with the present dataset, it is hard to tease apart the effects of confounding factors such as anthropogenic light, human presence and other urban factors, especially in field observations. Road noise, when specifically recorded and played back in controlled field or lab experiments, always had a noticeable effect (Schaub et al. 2008; Luo et al. 2015; Bunkley and Barber 2015; Finch et al. 2020; Vosbigian et al. 2024), but it was not the case when observed directly in the wild (Fig. 3b). This suggests that while acoustic stressors are potent, effects in complex, real-world environments can be masked or altered by confounding factors, highlighting the need for experimental approaches to isolate problematic acoustic cues for setting reliable management thresholds and informing engineering solutions that minimise harmful noise emissions. However, although data gaps remain in terms of species diversity, behavioural context (e.g. foraging, mating, migration), and regional representation, our findings support the development of group-level recommendations for mitigating ultrasonic noise pollution. Bats that use echolocation or communication above 20 kHz, which is true for most of the species, may face increased risks of noise exposure in future environments.

### Study locations

Investigations of anthropogenic noise on bats have been conducted predominantly in the Global North (Fig. 2a). Except for one study each from Brazil, Colombia, Mexico, Panama and South Africa (Geipel et al. 2019; Ramalho et al. 2021; Lara-Nuñez et al. 2022; Cory-Toussaint 2022; Yanten et al. 2022) and four studies in China (Haddock et al. 2019; Song et al. 2019, 2020; Wang et al. 2022), research is lacking in tropical regions, despite their significantly higher bat diversity. Roads and rails generally act as barriers to animal movement; therefore, it is particularly important to conduct more targeted studies on bats in tropical nations where human populations and transport networks significantly exceed those of most temperate regions.

### Affected species and research priorities in bat noise ecology

Almost 90% of bats use ultrasonic echolocation (>20 kHz) to forage and understand their environment (Altringham et al. 1996). However, the impact of noise does not always have to be associated with the behaviours linked with echolocation in bats. Acoustics also plays a role in prey detection, social communication, and other contexts requiring a broad understanding across species. Overall, more than 40 species are affected by traffic and/or anthropogenic noise, most from the species-rich Family Vespertilionidae. There is a need to shift focus to less-studied families that might also be vulnerable. False vampire bats (Family Megadermatidae) in Asia, Australia, and Central Africa remain unexplored but likely influenced by traffic noise. Megadermatids, often gleaning bats relying on prey-generated noise over echolocation, may be impacted similarly to Vespertilionid gleaners, which show sensitivity to traffic noise (Schaub et al. 2008; Siemers 2011; Bunkley and Barber 2015).

Whether this trend extends to other families of gleaning bats is unknown. Another understudied group is the non-echolocating Old-World Fruit bats (Family Pteropodidae). Although they do not use laryngeal echolocation, they are highly vocal (Christesen’ and Nelson 2000; Prat et al. 2017) and frequently urbanised, exposed to constant human disturbances and (traffic) noise. Yet, we found only one study that has investigated noise impacts on pteropodid vocalisation, finding a silentium effect in *Pteropus poliocephalus* (Pearson and Clarke 2019). This could hinder communication or reflect adaptive behaviour. Pteropodid bats, like *Pteropus*, use vocal cues in mother-pup interactions (Van Parijs and Corkeron 2002; Markus and Blackshaw 2002), and the effects of noise on such fine-scale communication in urban bat colonies warrant study. Vocal complexity and social communication are negatively influenced by traffic noise in the Asian particoloured bats (Haddock et al. 2019), but impacts on (audibly) highly vocal species such as flying foxes are unknown. However, anthropogenic stressors in urban-dwelling pteropodids such as *Pteropus medius* require particular attention because they are reservoir hosts of viruses implicated in recurring epidemiological outbreaks in India (Yadav et al. 2020; Sudeep et al. 2021), and they have been shown to be sensitive to human stressors such as increased light pollution (Murugavel et al. 2023).

### Acoustics of noise

We compared the results from studies that used either intensity or frequency or both to assess the impact of noise on bats. However, not all reported effects could be directly attributed to these acoustic parameters; most were correlated with noise presence rather than linked to a specific parameter. For example, bat activity was quantified near roads relative to other sites (Berthinussen and Altringham 2012; Bennett and Zurcher 2013; Cory-Toussaint 2022), and any observed effects were attributed to the road noise alongside other factors, but not exclusively to intensity or frequency when measured. Some studies reporting noise frequencies had either selected bands unlikely to overlap with bat calls or excluded certain frequencies from analyses (Domer et al. 2021; Cory-Toussaint 2022). Others described frequency information but did not explicitly incorporate it into result interpretation (Bunkley et al. 2015; Lehrer et al. 2021). Using playback experiments of music, Hooker et al. (2023) systematically tested music impacts while controlling for background vehicular noise, light pollution, and human activity, and reported reduced bat activity during music treatments. By contrast, Buxton et al. (2020) described low-frequency motorcycle rally noise affecting bats and other wildlife. In such cases, it is difficult to correlate specific acoustic parameters with reported impacts, underscoring the need for controlled experimental designs.

From the bat’s perspective, overlap between noise and call frequencies would be expected to cause direct impacts, highlighting the importance of studies comparing traffic noise across overlapping and non-overlapping ranges. Finch et al. (2020) conducted a Before-After Control Impact (BACI) experiment in the wild, exposing bats to both sonic and ultrasonic traffic noise levels, finding negative effects at both frequency ranges in free-flying bats. This suggests that low-frequency urban noise may also directly impact bats because their social calls are typically in this range, enabling communication over longer distances. For example, *Pipistrellus kuhlii* in urban habitats emitted social calls at lower frequencies than in less urban settings (Russo and Jones 1999), suggesting possible adaptive mechanisms unfolding in urban environments. However, more controlled studies focusing exclusively on acoustic parameters are essential to advancing our understanding of how traffic noise affects bats’ sensory ecology and functional roles in urban environments.

### Methodological concerns

The primary challenge in measuring the impacts of noise and its acoustic parameters on bats lies in how noise is quantified in these studies. Not all studies report both intensity and frequency information, and those that provide intensity information often fail to specify the distances at which measurements were taken. In some studies, noise is assessed subjectively, e.g., described as ‘loud’ or ‘quiet’ based on the observers’ perception (Zurcher et al. 2010; Yosef et al. 2021), or classified as ‘high’ or ‘low’ according to traffic volume, such as the number of vehicles passing in a given period (Myczko et al. 2017), with impacts discussed on that basis. While such approaches can offer valuable situational insights, they limit the extraction of precise acoustic parameters, making it difficult to place findings within a comparable framework for studying noise effects. A possible recommendation would be to adopt standardised methodologies, including measuring noise levels at defined distances, reporting the equipment used, and aligning measurements as closely as possible with existing literature to facilitate comparability.

### Beyond bats

Our findings, although centred on bats, highlight broader conservation concerns for other taxa with ultrasonic hearing capacity. The critical thresholds we identified ∼20 dB SPL intensity and ∼30 kHz frequency, above which masking effects become prevalent, are not unique to bats but overlap with the auditory sensitivity of other groups, including many nocturnal insects (Holderied et al. 2018; Geipel et al. 2021; Barber et al. 2022). For example, it is unclear how moths that are sensitive to ultrasound will respond to future soundscapes with heightened ultrasonic noise. Recent research has shown that anti-bat ultrasound production is widespread among moths, with an estimated 20% of Macroheterocera species producing ultrasonic responses to bat attacks (Barber et al. 2022), suggesting that interference in these frequency ranges could have broad ecological consequences for predator and prey species alike. Hence, EV-related ultrasonic emissions, if not further quantified and left unregulated, could pose risks to species that rely on high-frequency cues for predator avoidance (Corcoran and Conner 2012; Holderied et al. 2018). Similarly, ultrasonic communication, which can be crucial for social cohesion and reproductive behaviour, may be disrupted in other mammals, such as rodents or primates (Heffner and Heffner 1985; Brudzynski 2009; Ramsier et al. 2012; Merten et al. 2014). Canids, including domestic dogs, also have well-documented sensitivity to ultrasound, with upper hearing limits extending to at least 45 kHz and the ability to detect low-intensity ultrasonic sounds (Blackshaw et al. 1990). Importantly, despite this risk, no studies have systematically tested whether environmental ultrasonic noise from human-made sources interferes with or masks these signals of these or other animals in naturalistic settings, revealing an urgent need for targeted research. Our bat-centred thresholds offer an initial benchmark for broader biodiversity assessments, and we advocate for the extension of this work to other taxa that may be similarly vulnerable in emerging urban soundscapes.

### Future direction and recommendations

The difficulty in defining a single cut-off frequency for problematic ultrasonic noise in wildlife conservation mirrors challenges in human noise management (Leighton 2018). As Leighton (2018) shows, guideline bodies inadvertently extend maximum permissible levels below 20 kHz, and individual hearing variability complicates simple frequency cut-offs. Similarly, our synthesis highlights that bats’ diverse hearing sensitivities preclude a single upper frequency threshold that could reliably safeguard all species (Bohn et al. 2006; Geipel et al. 2021). Rather than advocating a fixed limit, we echo Leighton’s call for thresholds based on empirical data that reflect species-specific sensitivities and exposure contexts. For bats, this underscores the urgency of experiments focusing on EV noise emissions, including pedestrian alert sounds and ultrasonic distance sensors used, assessing masking potential, behavioural responses, and ecological consequences across relevant frequency ranges. In this context it is noteworthy that also unmanned aerial vehicles, ‘drones’ (UAVs), use ultrasonic sensors for navigation, landing and obstacle avoidance. Yet, evidence for these sensors directly interfering with ultrasonically sensitive wildlife remains limited, too, but remains plausible. The global drone market, however, is projected to experience substantial growth in the near future (Battsengel et al. 2020; Kapustina et al. 2021), with investment in UAVs is estimated at billions of dollars in developed countries, driven by applications in logistics, public services in urban spaces, as well as agriculture and forestry – another interface for a human-wildlife conflict (de Castro et al. 2021).

While 17.8 kHz may serve as a practical lower bound for the ultrasonic regime in human guidelines, it remains within the audible range for many bats and other taxa, but at the same time 17.8 kHz is below the frequencies that are typical of bat’s echolocation calls used in orientation and foraging, thereby showing some potential for a compromise between human demands and biodiversity needs. However, current human-centred limits for noise reduction risk underestimating ecological impacts, provoking a green-green dilemma where measures to promote environmental sustainability (such as EV adoption) could unintentionally conflict with the protection of ultrasound-sensitive biodiversity. This requires experimental work to identify biologically meaningful thresholds, as well as cross-disciplinary dialogue between sensory ecologists, engineers, and policymakers.

## Supporting information

Murugavel_etal_Supplementary_Material

Murugavel_etal_Supplementary_table_2

## Acknowledgements

We acknowledge the Indian Institute of Science Education and Research Mohali, for funding support to B.M. and infrastructural support for the study. We thank the library services of the Indian Institute of Science Education and Research, Thiruvananthapuram, for their assistance with the methodology. We thank Jonas Beer (Oldenburg) for help with the acoustic recordings of traffic sounds. O.L. is grateful for funding to the Sonderforschungsbereich (SFB) 1372 ‘Magnetoreception and Navigation in Vertebrates’ (project-ID395940726) by the Deutsche Forschungsgemeinschaft (DFG).

## Data availability

All literature data used for the review and meta-analysis are provided in the supplementary files.

## Author contributions

BM - Conceptualisation, Data curation, Formal analysis, Investigation, Methodology, Validation, Visualisation, Writing – original draft, Writing – review & editing

MJ - Investigation, Supervision, Validation, Writing – review & editing

OL - Conceptualisation, Data curation, Funding acquisition, Project administration, Supervision, Validation, Visualisation, Writing – review and editing

