## Supplementary material for "Sounds of silence: electric mobility promises a quieter soundscape for wildlife, but may challenge ultrasonically sensitive species globally": Murugavel_etal_Supplementary_Material

**Supplementary methods:**

*Field measurements*

Acoustic recordings of EVs and HEVs were made at several locations in the city of Oldenburg, Germany and Riga, Latvia, between March and April 2025, using a mobile bat detector (BATLOGGER M2, Elekon AG, Luzern, Switzerland) allowing us to record the full-spectrum between 10 to 192 kHz range. In Oldenburg, recordings of traffic were made during the daytime to avoid bat or insect ultrasound signals during measurements, and under zero to low wind conditions and clear weather. The detector was held perpendicular to the traffic lane. Focal vehicles were sampled opportunistically when a single vehicle passed, i.e. without specific choices of brands or models, but always only a single vehicle was present. In Riga, the ultrasound spectrum was monitored in the first half of the night; however, no bats were recorded during two consecutive nights of detector walks in the city centre. Temperatures were very low for bat activity in general, and particularly in the Baltic Region, this early in their activity season.

Details on the locations and conditions of ultrasonic car emissions recorded and used for illustrations in Figure 1: *Tesla Model X, BYD atto 3, Škoda Elroq* (Fig. 1a, g, j), 30 km/h zone, Posthalterweg, roundabout, Wechloy, Oldenburg, Germany (53,15977/8,16871), 04:00 to 06:00 PM, 2025-03-03. *VW ID.3* (Fig. 1d), 50 km/h zone, Ammerländer Heerstraße, city centre, Oldenburg, Germany (53,14166/8,19808), 12:50 to 01:10 PM, 2025-03-04. *Electric public bus* (Fig. 1f), 50 km/h zone, Stone Bridge, city centre, Riga, Latvia (56,9442/24,10056), 11:10 to 11:50 PM, 2025-04-25.

**Supplementary figures:**

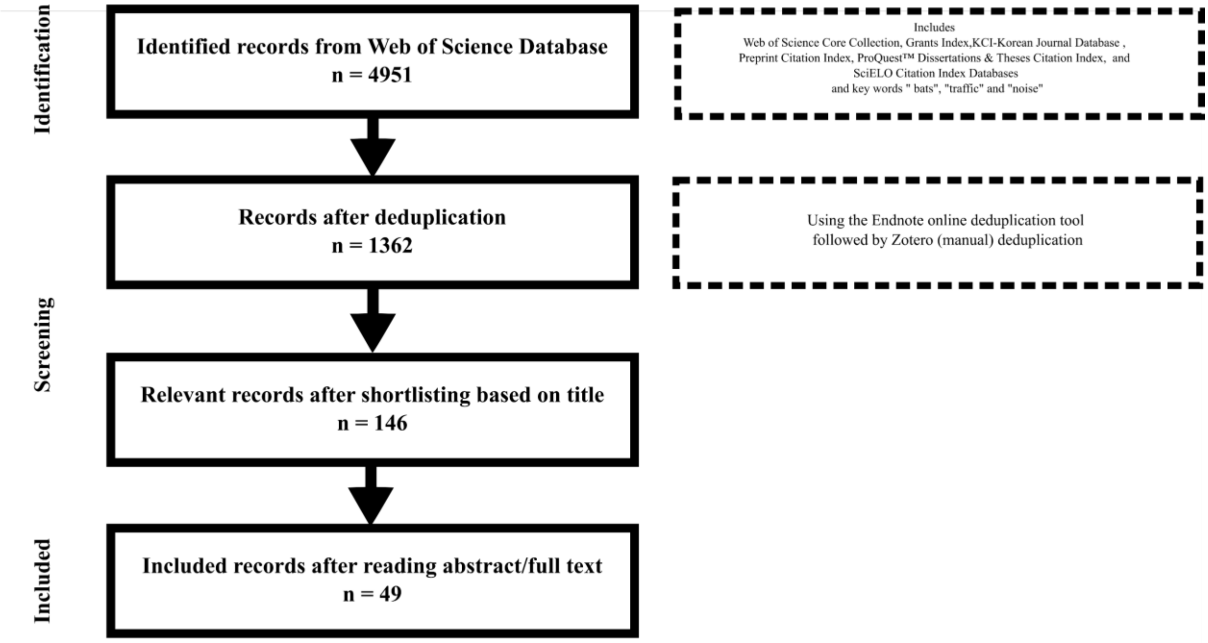


**Supplementary Fig.1.** Flowchart of literature survey following the PRISMA-EcoEvo guidelines. The entries are based on database search from the Web of Science databases

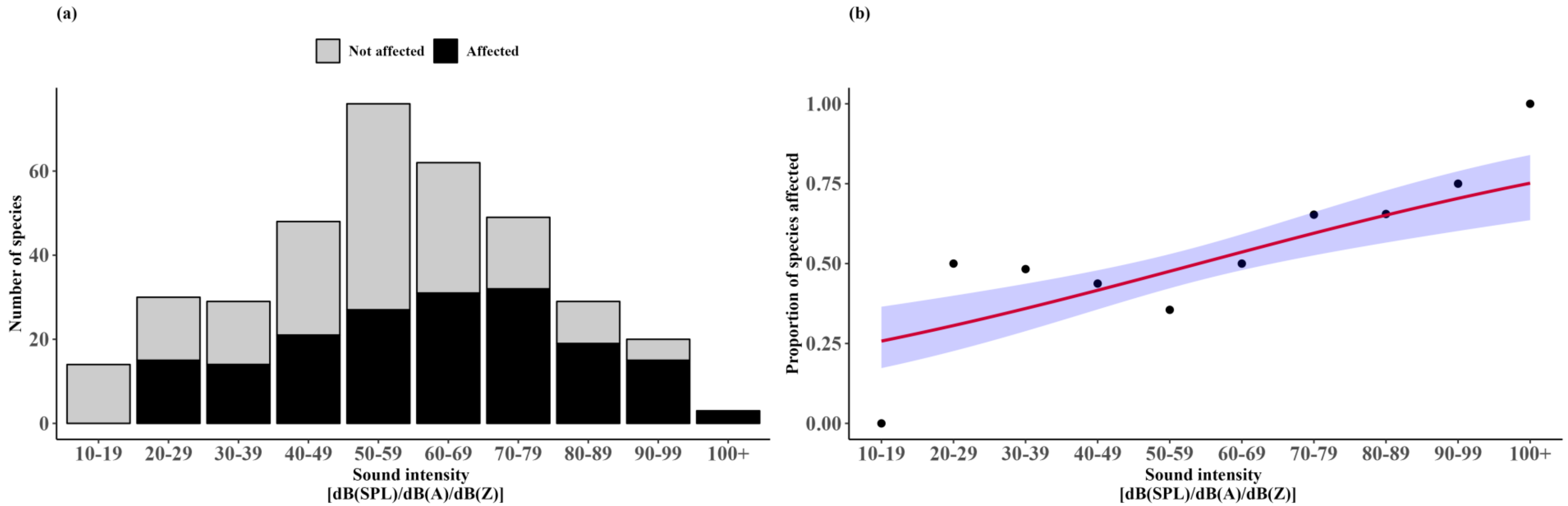


**Supplementary Fig.2.**(a) Number and (b) proportion of species/species groups exposed and affected by noise intensity categories. The predicted probabilities from the linear regression model are plotted as red lines with a 95% confidence (blue) ribbons.

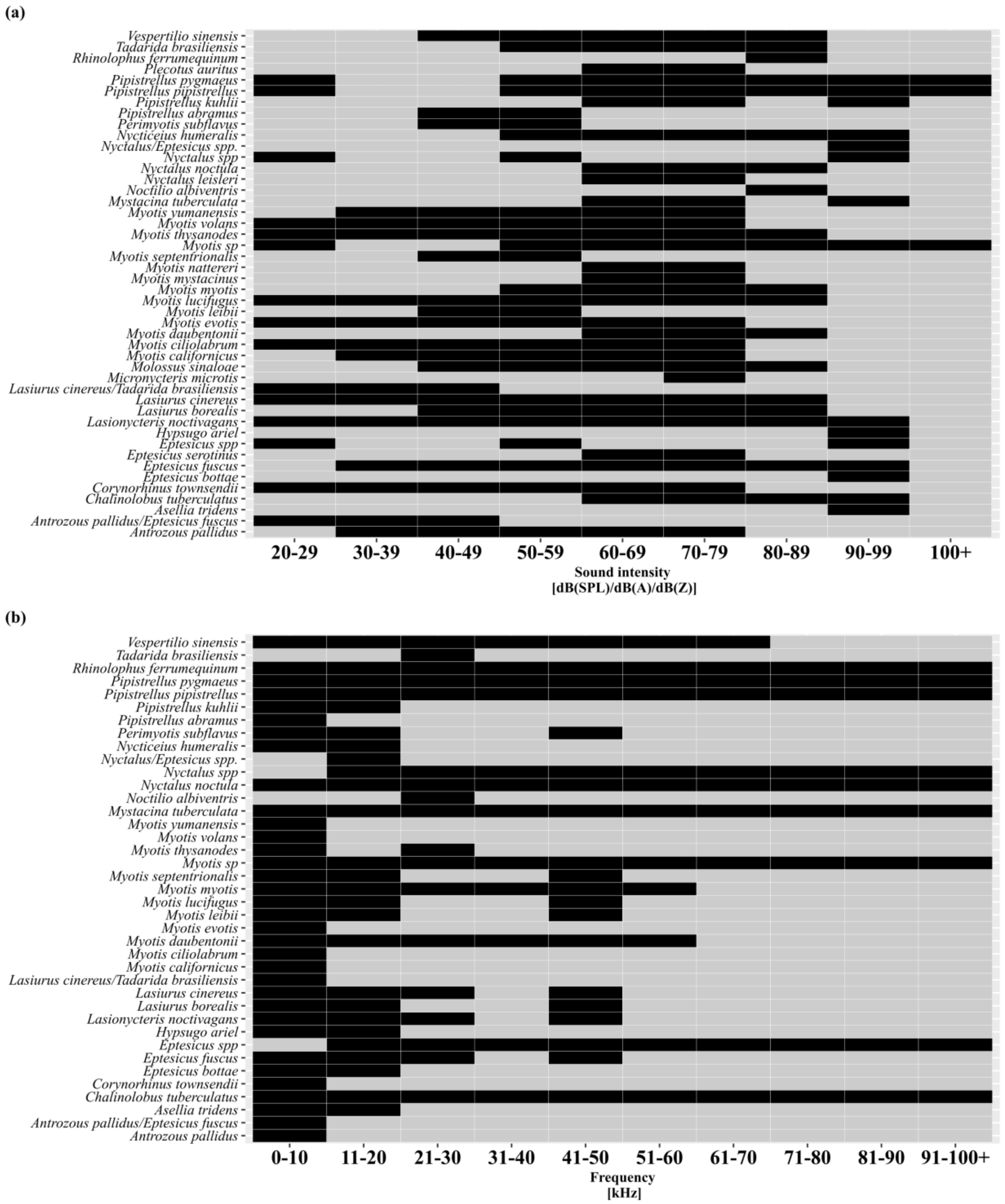


**Supplementary Fig.3.** Matrices showing the list of species that were affected either negatively or neutrally to the corresponding sound intensity (a) and frequency (b) categories.

**Supplementary Table 1. Number of studies and species/species groups investigated for noise types**

| Noise type | Number of Studies | Number of Species | Citation_List |
|---|---|---|---|
| Air | 6 | 20 | (Le Roux & Waas, 2012; Ednie, 2021; Jespersen *et al.*, 2022; Kuhlmann *et al.*, 2022; Wang *et al.*, 2022; Roswag *et al.*, 2025) |
| Other[+] | 15 | 76 | (Shirley *et al.*, 2001; Bunkley *et al.*, 2015; Rydell & Wickman, 2015; Warner, 2016; Pearson & Clarke, 2019; Gilmour *et al.*, 2020, 2021; Domer *et al.*, 2021; Gomes *et al.*, 2021; Lehrer *et al.*, 2021; Cory-Toussaint, 2022; Lara-Nuñez, Guerrero & Rizo-Aguilar, 2022; Brito *et al.*, 2023; Hooker *et al.*, 2023; Sereno-Cadierno *et al.*, 2025) |
| Rail | 2 | 6 | (Jerem & Mathews, 2021; Pakula & Furmankiewicz, 2022) |
| Road | 13 | 55 | (Kerth & Melber, 2009; Zurcher, Sparks & Bennett, 2010; Abbott, Harrison & Butler, 2012; Berthinussen & Altringham, 2012; Bennett & Zurcher, 2013; Bonsen, Law & Ramp, 2015; Myczko *et al.*, 2017; Borkin *et al.*, 2019; Laforge *et al.*, 2019; Medinas *et al.*, 2019; Bhardwaj *et al.*, 2020; Buxton *et al.*, 2020; Ramalho, Silveira & Aguiar, 2021) |
| Road* | 13 | 14 | (Schaub, Ostwald & Siemers, 2008; Siemers & Schaub, 2011; Luo *et al.*, 2014; Luo, Siemers & Koselj, 2015; Bunkley & Barber, 2015; Geipel *et al.*, 2019; Jiang *et al.*, 2019; Finch, Schofield & Mathews, 2020; Song *et al.*, 2020, 2020; Hart, 2022; Yanten, Cruz-Roa & Sanchez, 2022; Vosbigian *et al.*, 2024) |

[+] - indicates studies that used noise from Gas stations, compressor stations, music festivals, acoustic deterrents or general anthropogenic noises that could not be classified into any other category.

*-indicates studies that had used playback of road noise in either lab or field experiments.
